# Characterization of TR-107, a Novel Chemical Activator of the Human Mitochondrial Protease ClpP

**DOI:** 10.1101/2022.02.14.480426

**Authors:** Emily M. J. Fennell, Lucas J. Aponte-Collazo, Joshua D. Wynn, Kristina Drizyte-Miller, Elisa Leung, Yoshimi Endo Greer, Paul R. Graves, Andrew A. Iwanowicz, Hani Ashamalla, Ekhson Holmuhamedov, Henk Lang, Donald S. Karanewsky, Channing J. Der, Walid A. Houry, Stanley Lipkowitz, Edwin J. Iwanowicz, Lee M. Graves

## Abstract

We recently described the identification of a new class of small molecule activators of the mitochondrial protease ClpP. These compounds synthesized by Madera Therapeutics showed increased potency of cancer growth inhibition over the related compound ONC201. In this study, we describe chemical optimization and characterization of the next generation of highly potent and selective small molecule ClpP activators (TR compounds) and demonstrate their efficacy against breast cancer models *in vitro* and *in vivo*. One of these compounds (TR-107) with excellent potency, specificity and drug-like properties was selected for further evaluation. Examination of TR-107 effects in triple-negative breast cancer (TNBC) cell models showed growth inhibition in the low nanomolar range, equipotent to paclitaxel, in a ClpP-dependent manner. TR-107 reduced specific mitochondrial proteins including OXPHOS and TCA cycle components, in a time, dose and ClpP-dependent manner. Seahorse XF analysis and glucose deprivation experiments confirmed inactivation of OXPHOS and demonstrated an increased dependence on glycolysis following TR-107 exposure. The pharmacokinetic properties of TR-107 were compared to other known ClpP activators including ONC201 and ONC212. TR-107 displayed excellent exposure and serum t_1/2_ after oral administration. The antitumor response to TR-107 was investigated using human TNBC MDA-MB-231 cell line-induced xenograft tumors. Oral administration of TR-107 resulted in reduction in tumor volume and extension of survival in the treated compared with vehicle control mice. In summary, we describe the identification of highly potent new ClpP agonists with improved efficacy against TNBC, through targeted inactivation of OXPHOS and disruption of mitochondrial metabolism.

## Introduction

Triple-negative breast cancer (TNBC) is the most aggressive breast cancer subtype and is associated with poor prognosis, shorter progression-free and overall survival compared to other breast cancer subtypes(1–3). Unlike other subtypes, targeted therapies (e.g. tamoxifen and trastuzumab) are ineffective against TNBC leaving patients with limited options of systemic chemotherapy, surgery or radiation(3) or more recently immunotherapy(4,5). As an alternative, considerable interest has developed in targeting mitochondrial metabolism as a potential approach to treat recalcitrant cancers(6,7). Multiple studies now support the feasibility of targeting essential mitochondrial processes such as oxidative phosphorylation (OXPHOS), mitoribosomal activity and TCA cycle function as a unique strategy to inhibit cancer cell proliferation(8,9). This has led to the development of small molecule inhibitors of OXPHOS, TCA cycle enzymes (isocitrate dehydrogenases (IDH1/2), α-ketoglutarate dehydrogenase), as well as modifiers of cell death pathways(10,11). Included in this class of molecules are metformin and other clinical agents (BAY 87-2243, IACS-010759) designed to inhibit mitochondrial multimeric complex I enzyme that comprises the first step in the electron respiratory chain(12–14). While these have shown promising activities in preclinical studies, the clinical application of these agents has yet to be realized(8).

The small molecule imipridone ONC201 was initially identified in a screen for TRAIL inducers and later shown to have antagonistic effects on the dopamine receptors D2/D3(15,16). While the direct target was unresolved, multiple studies showed that ONC201 exhibited effective anti-cancer activities in diverse cancer models including breast, pancreatic, leukemia and others(15,17–23) leading to the advancement of ONC201 clinical evaluation.

In addition to an unresolved target, the chemical structure of ONC201 was initially reported incorrectly in a 1973 patent application of anticonvulsive agents, and was correctly reassigned decades later by Janda and colleagues(24). Furthermore, the unexpected finding that ONC201 potently affected OXPHOS and mitochondrial metabolism in models of TNBC suggested an alternative mechanism of action(25). This information, coupled with the correct chemical structure for ONC201, facilitated the recent discovery that ONC201 and chemically-related analogs (TR compounds) prepared by Madera Therapeutics, were activators of the mitochondrial matrix protease ClpP. This finding led to the initial identification of the mechanistic action of ONC201 and analogs as potential anti-cancer compounds(26,27).

Previously, modifiers of ClpP activity have demonstrated anti-cancer properties(28,29), with both inhibitors and activators of ClpP demonstrating anti-tumor efficacy for acute myelogenous leukemia(30,31). Additional chemical activators of ClpP include diverse chemical structures such as the macrocyclic acyldepsipeptides (ADEPs) and D9 (Fig S1A). ADEP and D9 were both described in 2018 as human ClpP activating agents prior to the discovery of a similar mechanism of action for ONC201 and the TR compounds(29). Madera was the first to prepare a number of highly potent, cell permeable ClpP activators through modifications of peripheral functionality on the chemical core of ONC201, and later through changes to the chemical core. The in depth understanding of the structure-activity relationship for these agents allowed the effective coupling of TR molecules (TR-79, TR-80) to agarose beads, thereby facilitating the identification of ClpP as the binding partner(26). Additional genetic *CLPP* knockdown studies confirmed ClpP as the biological target for ONC201, and a crystal structure of ClpP and ONC201 further validated ClpP as a direct and specific binding partner(27).

ClpP is a component of the ClpXP protein complex localized in the mitochondrial matrix. ClpP is a tetradecameric serine protease that forms a complex with hexameric AAA+ ClpX, an ATP-dependent protein unfoldase that enables substrate recognition and unfolding prior to its degradation by ClpP(31–34). The crystal structure of ClpXP has been solved from a number of species providing detailed insight into how this proteolytic protein complex is regulated. ClpX binds to a hydrophobic pocket in ClpP thereby facilitating the opening of the axial pore and passage of unfolded proteins into the central barrel of ClpP. Pharmacological activators of ClpP, such as the ADEPs, were the first small molecules shown to bind to this hydrophobic pocket and open the axial pore of ClpP in the absence of ClpX, allowing for nonspecific entry of proteins into the active site of ClpP(28,35,36). As part of the mitochondrial unfolded protein response, ClpXP canonically targets misfolded proteins to prevent formation of protein aggregates in the mitochondria(33,37). ClpXP also has regulatory roles in heme biosynthesis(38–40), mitophagy(41,42), and reduction in reactive oxygen species levels following mitochondrial depolarization(34). ClpP is overexpressed in breast cancer(43), providing the potential for utilizing highly selective and potent ClpP activators as a novel approach to disrupt mitochondrial metabolic processes required for TNBC proliferation.

In this study, we present data on the characterization of a novel class of highly potent and selective ClpP activators. We show that one, TR-107 induces the downregulation of select mitochondrial proteins, impairs OXPHOS and inhibits TNBC growth in a ClpP-dependent manner, with markedly improved potency compared to the imipridones (ONC201, 206, 212). Analysis of the pharmacokinetic properties of TR-107 demonstrated high systemic drug levels following oral administration. Furthermore, the efficacy of TR-107 in mouse xenograft models of MDA-MB-231 TNBC cells showed that TR-107 was well tolerated, and inhibited tumor growth and increased animal survival. Finally, we show that the anti-proliferative activity of ClpP-activating compounds is associated with inhibition of OXPHOS and loss of mitochondrial proteins involved in metabolism.

## Materials and Methods

### Cell Culture

Human TNBC cell line SUM159 was a generous gift from Dr. Gary Johnson (University of North Carolina at Chapel Hill). MDA-MB-231 (WT and CLPP-KO) cells were provided by Dr. Stanley Lipkowitz (National Cancer Institute). MDA-MB-231 cells were cultured in RPMI 1640 media (Gibco, 11875-093) supplemented with 10% fetal bovine serum (FBS; VWR-Seradigm, 97068-085), and 1% antibiotic–antimycotic (anti/anti; ThermoFisher Scientific, 15240062). SUM159 cells were cultured in Dulbecco’s modified Eagle’s medium: Nutrient Mixture F-12 (DMEM/F12; Gibco, 11330-032) supplemented with 5% FBS, 5 μg/mL insulin (Gibco, 12585014), 1 μg/mL hydrocortisone, and 1% anti/anti. Cells were maintained at 5% CO_2_ and 37°C and periodically tested for mycoplasma.

### CRISPRi cell lines

sgRNA against human ClpP (Eurofins Genomics) was annealed and ligated into vector VDB783. This vector was transformed into DH5α cells and the presence of sgRNA was confirmed by colony PCR. Lentivirus was produced in HEK293T cells following transfection with plasmid and jetPRIME (PolyPlus) as previously described(44). Clonal populations of SUM159 cells infected with dCas9-KRAB lentivirus were clonally isolated to ensure dCas9-KRAB expression. Isolated SUM159 cells were infected with ClpP sgRNA lentivirus followed by additional selection passage in growth media supplemented with 2.5 mg/ml puromycin.

### Synthesis of Compounds

All TR compounds were prepared by Madera Therapeutics, LLC as previously described in the patent literature(45). ONC201, ONC206, and ONC212/TR-31 were purchased from Selleck Chemicals, LLC (S7963, S6853, S8673, respectively).

### Viability Assays

Total cell counting assays were performed by seeding MDA-MB-231 cells (WT and ClpP-KO; 4000 cells/well) or SUM159 (WT and ClpP-KO; 1000 cells/well) in a 96-well plate (Perkin Elmer, 6005050) and allowed to adhere overnight. Growth medium was then aspirated and replaced with 100 μL of growth medium supplemented with drug at concentrations indicated in the figure or figure legends. Cells treated with vehicle control (0.1% DMSO, Sigma-Aldrich D2650) were used as a negative control in all experiments and cells were incubated with drug-containing media for indicated timepoints. Media was then aspirated and replaced with 100 μL DPBS (Gibco, 14190-144) containing 1 μg/mL Hoechst stain (ThermoFischer Scientific, H3570) and allowed to incubate at 37°C for 30 minutes. Total cell number was then quantified using the Celigo Imaging Cytometer (Nexcelom).

### Immunoblotting

MDA-MB-231 or SUM159 cells were plated (100,000 cells/well) on a 6-well Costar plate (Corning, 3516) and incubated with compounds as described above for cell viability assays. Following treatment, cells were rinsed three times with 2 mL of cold DPBS and lysed using RIPA buffer (20 mM Tris (pH 7.4), 137 mM NaCl, 10% glycerol, 1% Nonidet P-40, 0.5% deoxycholate, 2 mM EDTA) supplemented with 2 mM Na3VO4, 10 mM NaF, 0.0125 μM calyculin A, and complete protease inhibitor cocktail (Roche Diagnostics, 11873580001). Cell lysates were clarified and immunoblotted as described earlier(46). Membranes were incubated with primary antibodies (Supplemental Information) diluted 1:1000 in 1% fish gelatin (Sigma Aldrich, G7041)/TBST overnight at 4°C, removed, washed 3 × 5 minutes in TBST, and placed in their respective 2° antibodies (1:10,000 dilution in 5% milk/TBST) for 1 hour. Membranes washed 3 × 5 minutes in TBST then incubated in ECL reagent (BioRad, 170-5061) and imaged using a Chemidoc MP (BioRad). Acquired images were processed/quantified using Image Lab software (BioRad).

### Caspase Activity Assay

Caspase-3/7 activity was analyzed using a fluorescent peptide substrate (Ac-DEVD-AMC, Cayman Chemical, 14986) as previously described(46). Briefly, cells were plated at 1 × 10^5^ cells per well (6 cm^2^ plate, Corning, 430166) and treated for 24 hours with 10 μM ONC201, 150 nM TR-57, 100 nM TR-107 (10X IC50) as well as 0.1% DMSO or 100 nM staurosporine. The samples were collected in lysis buffer (50 mM HEPES (pH: 7.4), 5 mM CHAPS, and 5 mM DTT), and caspase activity was measured by monitoring fluorescence (ex/em: 360 nm/ 460 nm) using a SpectraMax i3x (Molecular Devices).

### ClpP Activity Assay

Measurement of *in vitro* activity of purified recombinant human ClpP was performed with a slight modification to the method previously described(47). Briefly, ClpP activity was measured through degradation of casein-FITC (Sigma-Aldrich, C0528) in the presence of indicated compounds. The activity of ClpP proteolytic subunit (41 μg/mL(1.4 μM)) was measured in assay buffer (20 mM HEPES (pH 8.0), 100 mM KCl, 1 mM DTT, and 100 μM ATP) using 235 μg/mL fluorogenic casein-FITC substrate. Enzyme and compounds were mixed and incubated in assay buffer for 30 minutes before adding casein-FITC substrate. The kinetics of substrate degradation fluorescence was monitored in 96-well plates (PerkinElmer, 8059-21401), and fluorescence was recorded at 490 nm/ 525 nm (ex/em) using SpectraMax i3x (Molecular Devices). Linear slope of fluorescence signal was used to measure ClpP activity and normalized to DMSO control, expressed as RFU/hour.

### Surface Plasmon Resonance (SPR)

For the SPR experiments, recombinant human ClpP was purified as previously described(47). SPR measurements were performed on a Biacore X100 instrument (Cytiva Life Sciences, Marlborough, MA, USA) at 25°C in buffer (25 mM sodium phosphate, pH 7.5, 200 mM KCl, 0.05% Tween-20, and 0.004% DMSO). ClpP was immobilized onto flowcell 2 of a CM5 chip (Cytive Life Sciences, Marlborough, MA, USA) using the amine coupling wizard using Biacore X100 Control Software. A pulse of a mixture of NHS and EDC activated the CM5 chip surface. ClpP was diluted to 100 μg/ml in 10 mM sodium acetate, pH 4.5 immediately prior to use and injected over the activated surface for 180 seconds. Remaining activated sites were blocked with a pulse of ethanolamine. 12000 RU of ClpP was captured using this procedure. Flowcell 1 was unmodified and served as a reference. For TR-107 binding to ClpP, a concentration series of TR-107 in DMSO was diluted into buffer to achieve 0.004% final DMSO concentration in all samples. Three replicates were injected for each concentration. Data were blank subtracted and analysis was performed in Biacore X100 Evaluation Software (Cytiva Life Sciences, Marlborough, MA, USA). The steady state response was calculated at 15 seconds before the end of injection with a window of 15 seconds. Data were fitted to a one-site Langmuir binding model.

### Mitochondrial Respiration Analysis

Cellular oxygen consumption and extracellular acidification rates (OCR and ECAR, respectively) were measured using Seahorse XFe96 (Agilent, Santa Clara, CA, USA). MDA-MB-231 cells were plated (15,000 cells/well) in Seahorse XF96 plates (Agilent, 102416-100) and allowed to adhere overnight. Cells were then treated with 25 or 50 nM TR-107 for 24 hours. On the day of Seahorse XF analysis, growth medium was replaced with XF base medium (Agilent, 103575-100) supplemented with appropriate concentrations of TR-107. After measurement of basal OCR/ECAR, 1 μM Oligomycin, 1 μM FCCP, 1 μM Rotenone/Antimycin A (Agilent, 103015-100) were added at indicated timepoints. OCR/ECAR were measured every 6.5 minutes for 73 minutes.

### Mitochondrial DNA (mtDNA) copy number analysis

Mitochondrial DNA copy number was determined as previously described(25). Briefly, genomic DNA was isolated from tissue lysate with DNeasy Blood & Tissue kit (Qiagen, 69504). mtDNA copy number was determined by quantitative PCR with Human mitochondrial to nuclear DNA ratio kit (Takara Bio USA, 7246).

### Pharmacokinetic Analysis

The compounds were evaluated for pharmacokinetic properties in ICR mice (Sino-British SIPPR/BK Lab Animal Ltd, Shanghai, China) by Shanghai Medicilon, Inc. The compounds were administered either intravenously (tail vein) or orally (oral gavage) in vehicle (5% DMSO, 10% Solutol [Kolliphor HS 15, Sigma Aldrich, 42966] in sterile DPBS, 85% water) to three male mice per study arm. Blood was taken via the orbital venous plexus (0.03 mL/timepoint) at 0.083, 0.25, 0.5, 1, 2, 4, 8 and 24 hours, unless otherwise noted. Samples were placed in tubes containing heparin sodium and stored on ice until centrifuged. Blood samples were centrifuged at 6800 x g for 6 minutes at 2-8°C within 1 hour following sample collection and stored at −80°C. Proteins from plasma sample aliquots (20 μL) were precipitated by the addition of MeOH (400 μL) containing an internal standard (tolbutamide,100 ng/mL), vortexed for 1 minute, and centrifuged at 18,000g for 7 minutes. Supernatant (200 μL) was transferred to 96 well plates for analysis.

Analysis of samples was conducted as follows. An aliquot of supernatant (1 μL) was analyzed for parent compound by LC-MS/MS using a Luna Omega C18 column (2.1 × 50 mm, 1.6 μm; Phenomenex, 00B-4747-B0) with 0.1% formic acid/water and 0.1% formic acid/acetonitrile gradient system, and a TQ6500+ triple quad mass spectrometer (positive ionization mode). In the case of TR-107, the parent compound and internal standard (IS) were detected with electrospray ionization in positive mode (ESI+) using multiple-reaction monitoring (MRM) of mass transition pairs at m/z of 391.2/247.2 (TR-107) and 271.1/172.0 (IS, tolbutamide) amu. The calibration curve was obtained by spiking known concentrations (2 to 2000 ng/mL) of TR-107 into blank mouse plasma. The PK parameters including Area Under the Curve (AUC_(0-infinity)_) elimination half-life (t_1/2_), maximal plasma concentration (C_max_), and oral bioavailability (F%) were analyzed by noncompartmental methods.

### Protein Binding Studies to Plasma Proteins

Protein binding studies were performed by Eurofins Panlabs, Inc. This assay utilizes equilibrium dialysis in a microplate format, as previously described(48). Briefly, this analysis is used to determine the bound and unbound fraction of the drug and calculate the percentage of the test compound binding to murine plasma proteins.

### Mouse Xenograft Studies

All mouse xenograft studies were performed at Charles River Laboratories (Morrisville, NC) using their established procedures. Detailed methods are described in Supplemental Methods. In brief, CR female NCr nu/nu mice were injected with 5 × 10^6^ MDA-MB-231 tumor cells suspended in Matrigel orthotopically into the flank mammary fat pad. Cell injection volume was 0.05 mL/mouse. A pair match was performed when tumors reached an average size of 60-100 mm^3^ (8-12 weeks) after which treatment with TR-107 was initiated. TR-107 (Madera Therapeutics-Lot No. 3 and 4) was dissolved in vehicle (5% DMSO, ~5% Solutol [Kolliphor HS 15] in sterile DPBS). Each week appropriate amounts of TR-107 were dissolved in DMSO and these solutions were further diluted with prefiltered 5% Solutol in DPBS to yield dosing solutions at 0.4 and 0.8 mg/mL, which provided 4 and 8 mg/kg, respectively, in a dosing volume of 10 mL/kg, adjusted for body weight. Dosing solutions were stored at 4°C. Tumor size and volume were measured and calculated as described in Supplemental Information, as was body weight and animal health.

### Statistical Analysis

Statistical analysis for viability, mitochondrial respiration, ClpP activity assays, and animal studies was performed using Prism 9 (GraphPad, San Diego, CA, USA). Analysis of pharmacokinetic data was performed using FDA certified pharmacokinetic program Phoenix WinNonlin 7.0 (Pharsight, Mountain View, CA, USA).

## Results

### New TR chemical scaffolds are highly potent inhibitors of TNBC growth

Madera Therapeutics synthesized multiple small molecules containing modifications to the multi-ring core system using the optimized peripheral functionality from TR-65 and similar analogs. This included modification of the core imipridone structure with the appropriate addition of nitrile, halide and trifluoromethyl groups to the two benzyl moieties. The structures of these compounds and other established ClpP agonists (i.e. ADEPS and D9) are shown in Fig. 1A, Fig. S1A, or a previous publication(26).

**Figure 1.**
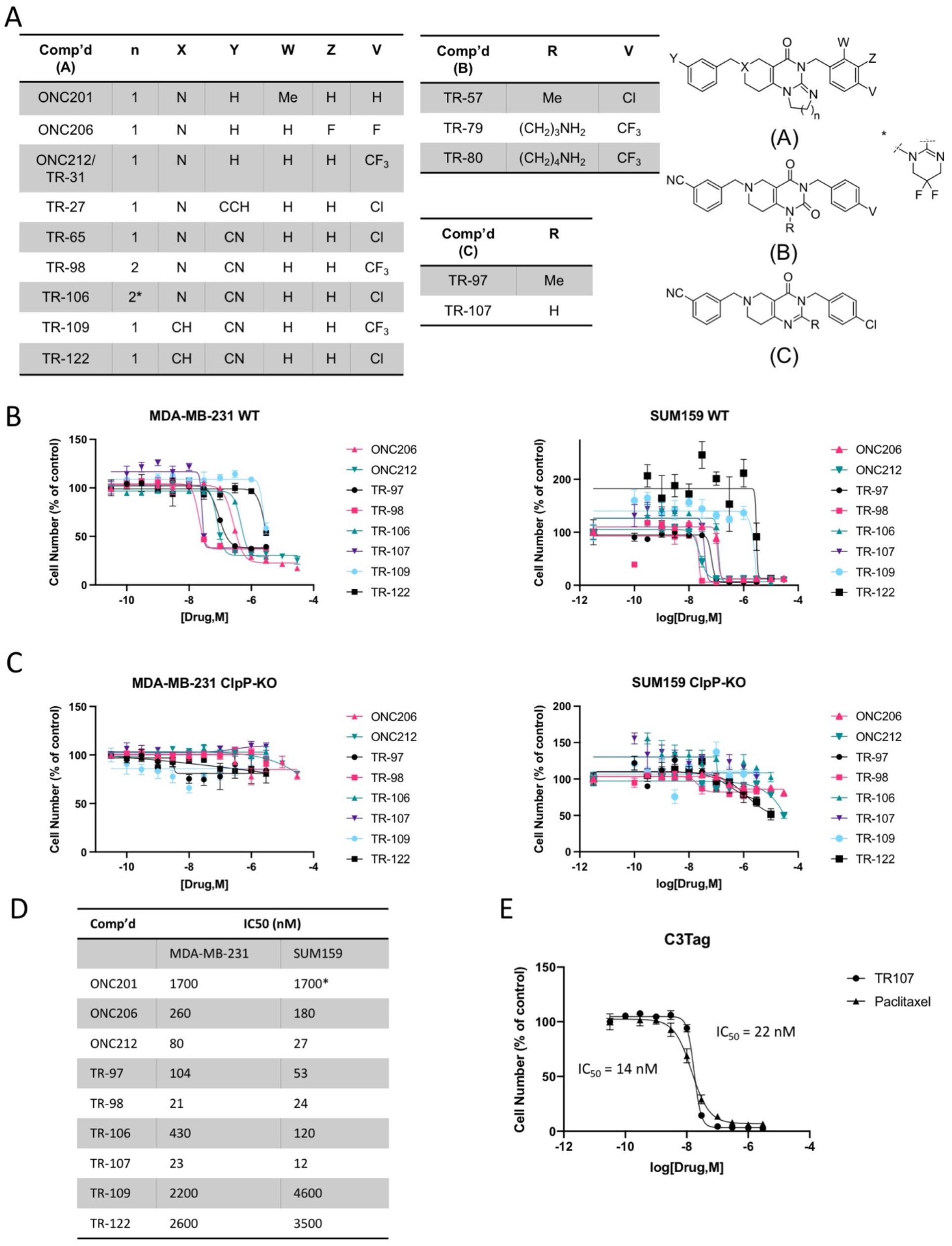
New TR compound analogs potently inhibit breast cancer cell growth in a ClpP-dependent manner. A) Comparison of novel TR compound chemical structures to ONC201 and analogs. Cell viability assays of (B) Wildtype (WT) and (C) ClpP knockout (ClpP-KO) MDA-MB-231 and SUM159 cells. Cells were treated with indicated drug concentrations for 72 hours and imaged following addition of Hoechst stain as described in Materials and Methods. Values represent mean ± SEM normalized to DMSO control, representative of N=3. D) IC_50_ values in WT cells for compounds represented in panel B. Values represent mean IC_50_ value (nM), N=3. *ONC201 IC_50_ in MDA-MB-231 obtained from data shown in Fig. 4D, SUM159 cells previously published^25^. E) Cell viability assay of C3Tag cells following TR-107 and paclitaxel treatment. Cells were treated with drug concentrations as indicated for 72 hours and imaged following Hoechst stain addition. Representative of N=2. Average IC_50_ values indicated to the left and right of the graph for paclitaxel and TR-107, respectively.

We tested the TR compounds’ ability to inhibit the growth of two commonly studied TNBC cell lines. Many of these new compounds showed significantly enhanced potency of cell growth inhibition in the MDA-MB-231 and SUM159 cell models compared to ONC201 and ONC206. Of these, TR-98 and TR-107 demonstrated the greatest growth inhibitory potency, comparable to the previously reported TR-57 and TR-65(26). For example, TR-107 inhibited cell growth with an IC_50_ of 12 nM and 23 nM in SUM159 and MDA-MB-231 cells, respectively (Fig. 1B). By contrast, TR-109 and TR-122, devoid of a key nitrogen atom of the 6 membered 3-piperideine system (X in structure A, Fig. 1A), were less potent, suggesting the importance of a reported hydrogen bond interaction with tyrosine 118 of ClpP(27). This hydrogen bond is one of three hydrogen bonds observed in the human ClpP-ONC201 complex structure, involving glutamate 82 and glutamine 107 (through a bridging water molecule) that anchor the molecule to the hydrophobic site of ClpP(27). Taken together, our data with TR-97, TR-98 and TR-107 show that significant chemical modification of the imipridone core is tolerated and that the imidazoline moiety of the imipridone core is not required for high potency.

To confirm the ClpP dependence of the new TR compounds, the growth inhibitory effects of these compounds were compared in matched pairs of wild-type (WT) and ClpP-KO MDA-MB-231 and SUM159 cell lines. Compared to the dose-dependent inhibition observed in the WT cells, no significant growth inhibition was observed in the ClpP-KO cells, providing evidence that ClpP is the major target for these molecules (Fig. 1C). To compare the response to TR-107 in another model of TNBC, we examined the effects of TR-107 on C3(1)-Tag cells and compared the potency of growth inhibition to the tubulin inhibitor paclitaxel, an established therapeutic. The C3(1)-Tag model is a transgenic murine model of TNBC that shares many of the essential characteristics of human TNBC(49). TR-107 potently inhibited C3(1)-Tag cell proliferation with an IC_50_ value (22 nM), similar to paclitaxel (14 nM) (Fig. 1D), which was also similar in potency to that observed with the MD-MBA-231 and SUM159 cells (23 and 12 nM, respectively).

### Characterization of novel activators of ClpP

We examined the ability of the new TR compounds to bind and activate ClpP. Select TR compounds were incubated with purified recombinant human ClpP and ClpP proteolytic activity was determined using the *in vitro* activity assay described in Materials and Methods. All compounds showed dose-dependent increases in ClpP activity with the greatest potency of ClpP activation observed with TR-107, followed by TR-98 and TR-97 (EC50 = 140, 310, and 340 nM respectively). Consistent with reduced effects on cell growth, TR-109 and TR-122 were also weaker activators of ClpP *in vitro* (Fig. 2A). Direct binding of TR-107 to purified ClpP was also examined. Surface plasmon resonance (SPR) measurement of TR-107 binding to ClpP, revealed values equivalent to that observed for ClpP activation (K_d_ ~180 nM) (Fig. 2B). Higher concentrations of compound were required for ClpP activation or binding *in vitro*, compared to cell growth inhibition. However, compounds that potently activate ClpP *in vitro* also potently inhibited cell growth as observed above (Fig. 1) and as previously reported(26).

**Figure 2.**
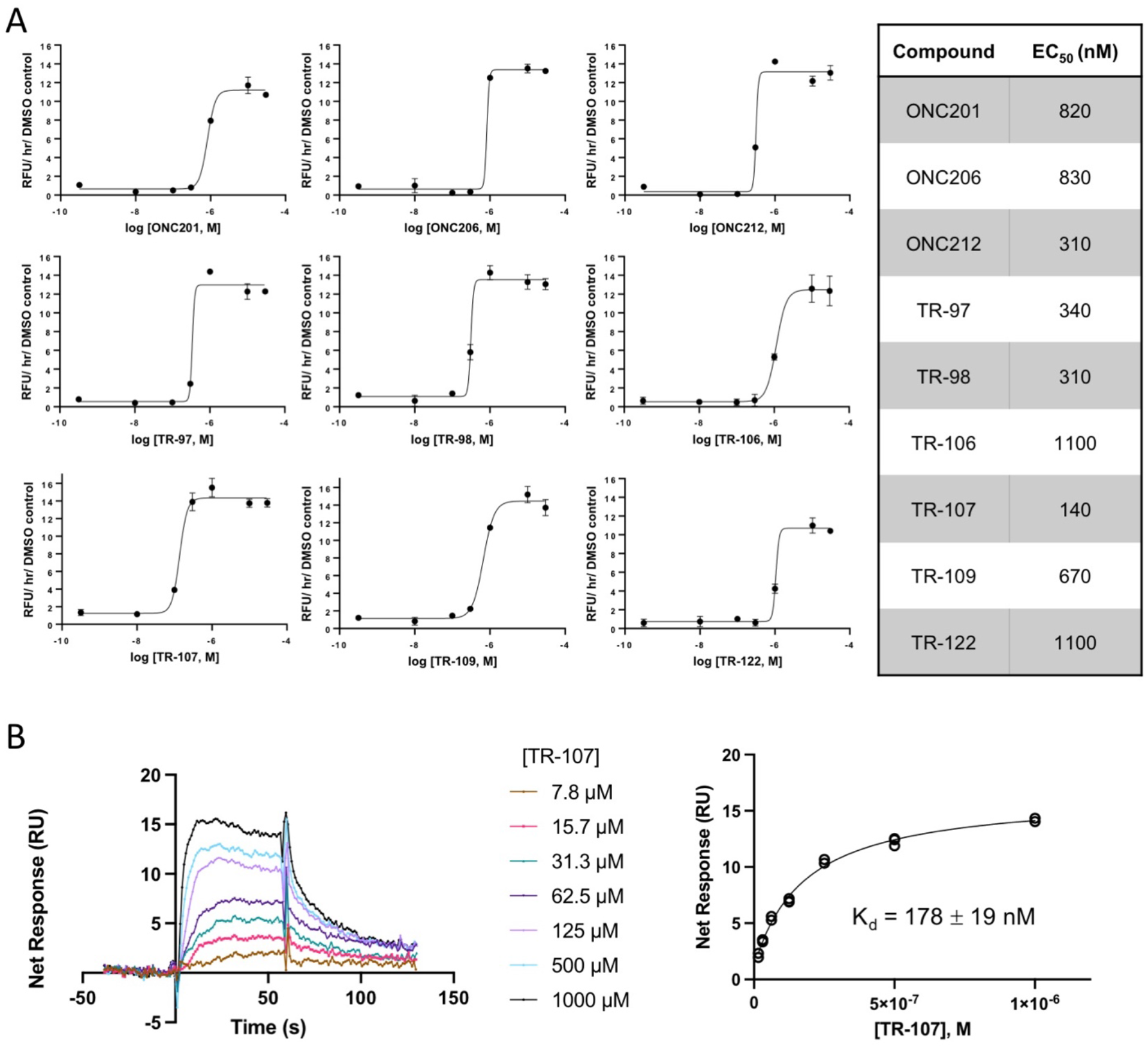
New TR compounds are potent activators of mitochondrial ClpP. A) *In vitro* ClpP activity assays was performed as described in Materials and Methods using fluorescent casein-FITC. Increase in fluorescent intensity relative to DMSO controls is shown; values represent mean ± SEM for technical replicates. EC_50_ values are presented in the table to the right. B) Left panel-surface plasmon resonance (SPR) sensorgrams of TR-107 binding to recombinant human ClpP coupled on the chip. Right panel - binding curves are shown as response units (RU) vs. ligand concentration at steady state and fitted to a one-site Langmuir binding model. The apparent K_d_ value obtained from the fit is shown.

### TR-107 inhibits TNBC growth without a significant increase in apoptosis

Since TR-107 showed excellent potency in these assays, this compound was selected for further testing on cell growth inhibition and apoptosis. MDA-MB-231 or SUM159 cells were incubated with TR-107 and total cell counts determined every 24 hours for up to 72 hours following treatment at the indicated concentrations. As shown in Fig. S1B, TR-107 potently (at concentrations >10 nM), inhibited the increase in cell number compared to vehicle control in both cell lines. By comparison, cell growth was not inhibited in the ClpP-KO cells with TR-107 concentrations as high as 1 μM (Fig. S1B). Because the highest concentration of TR-107 tested in MDA-MB-231 or SUM159 cells, did not reduce the total cell number below the initial seeded number, we tested for evidence of apoptosis.

Caspase-3/7 assays were performed on lysates from the MDA-MB-231 or SUM159 cells following 24-hour compound treatment at indicated concentrations. Caspase activity was quantified as described in Materials and Methods. Staurosporine (STS)-treated cells showed a strong increase in caspase-3/7 activity as expected, whereas no increase in caspase-3/7 activity was detected in lysates from ONC201, TR-57, or TR-107 treated cells (Fig. S1C). Similarly, no increase in PARP cleavage after ONC201, TR-57 or TR-107 treatment was observed (data not shown). Taken together, these findings indicate that the TR compounds show growth inhibitory effects without substantially increasing apoptosis, consistent with previously reported results for ONC201 in MDA-MB-231 cells(25).

### TR-107 induces time- and dose-dependent reduction of multiple mitochondrial proteins

We and others reported that treatment of MDA-MB-231 or SUM159 cells with ONC201 or TR-57 resulted in the reduction of the mitochondrial proteins elongation factor Tu (TUFM) and transcription factor A (TFAM)(25,26). Since ClpP is a mitochondrial matrix protease, we compared the time and dose-dependent effects of TR-107, TR-57 and ONC201 on additional mitochondrial proteins in MDA-MB-231 and SUM159 cells. As determined by immunoblotting, both TUFM and TFAM protein levels began to decline ~6 hours after treatment with 100 nM TR-107, 150 nM TR-57 or 10 μM ONC201 and showed a complete or near-complete loss by 24 hours (Fig. 3A; Fig. S2A, C). We also observed the time- and dose-dependent loss of aconitase (ACO2) and isocitrate dehydrogenase (IDH2), two key TCA cycle enzymes beginning at ~12-24 hours following treatment (Fig. 3A-B; Fig. S2A-D). Succinate dehydrogenase A (SDHA) and complex I subunit NDUFS3, both OXPHOS proteins, were observed to decline by 12 to 24 hours (Fig. 3A, 3C; Fig. S2A, S2C). ClpX, the ATPase subunit of the ClpXP complex, was previously observed to decline after 24 hours ONC212 treatment in pancreatic cancer models(50). We observed ClpX to decline rapidly (~3 hours) after TR compound addition (Fig. 3A; Fig. S2A). The loss of all these proteins was dose-dependent and occurred at concentrations at or above the respective growth inhibitory (IC_50_) value for each compound with TR-107 showing the most potent effects (Fig. 3B; Fig. S2B). By comparison, the levels of these proteins were not affected in the ClpP-KO cell lines, even at the highest concentration of compound tested (Fig. 3, Fig. S2).

**Figure 3.**
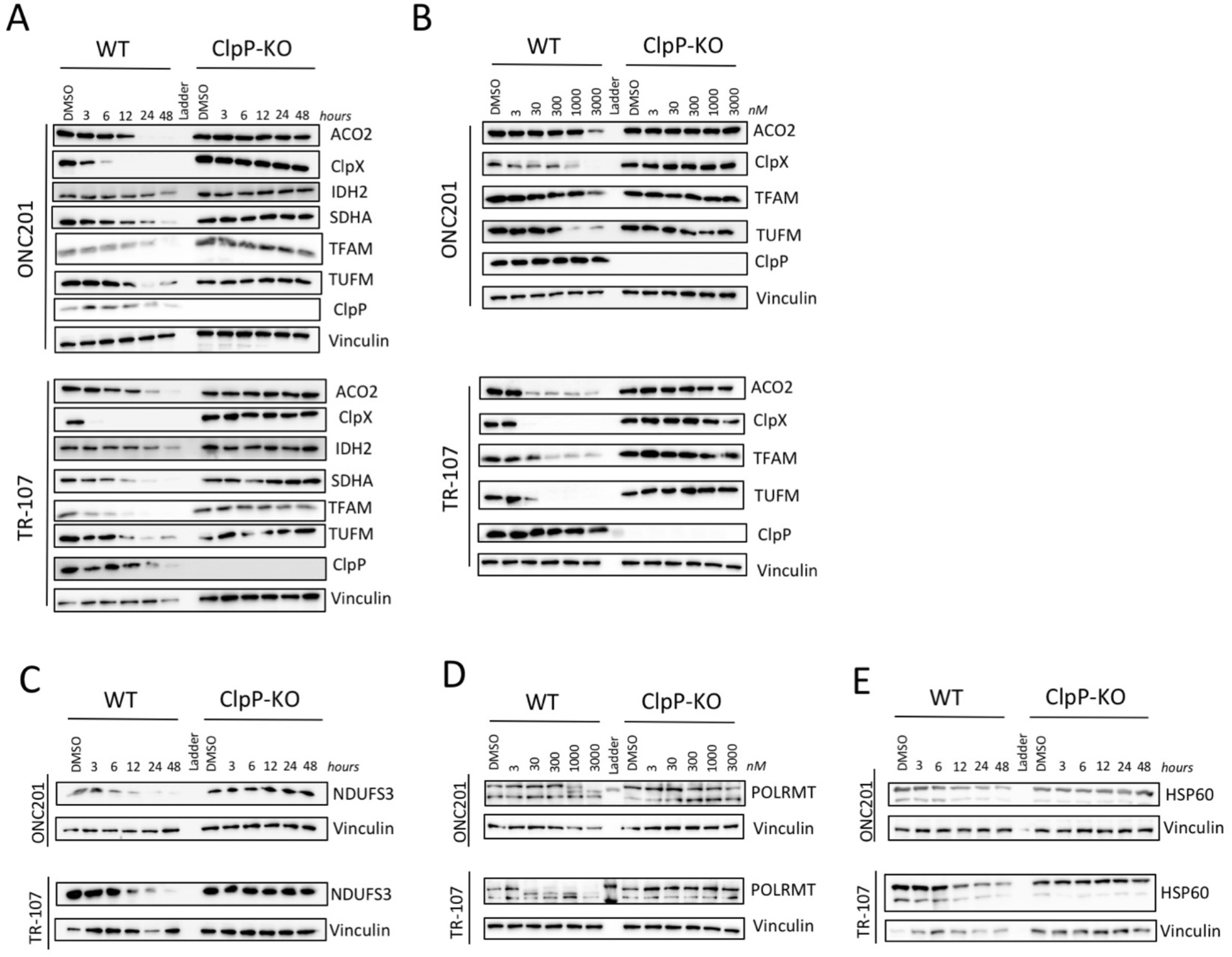
TR-107 induces loss of mitochondrial proteins in MDA-MB-231 cells in a ClpP-dependent manner. MDA-MB-231 cells (WT or ClpP-KO) were treated with 10 μM ONC201 or 100 nM TR-107 for indicated timepoints or with indicated doses of ONC201 or TR-107 for 24 hours. Immunoblots were performed for (A, B) mitochondrial metabolic proteins and ClpX, (C) NDUFS3, (D) POLRMT, and (E) HSP60 as described in Materials and Methods. Images are representative of N=3 (A,B) or N=2 (C-E) blots.

DNA-directed RNA polymerase, mitochondrial (POLRMT) is a TFAM interacting protein involved in the replication of mitochondrial DNA. Recent studies have shown the efficacy of inhibiting POLRMT as a potential anti-cancer approach(10). Because of the observed decline in TFAM, we investigated whether POLRMT was similarly affected by these treatments. Immunoblotting data showed that POLRMT followed a dose-dependent decline in full length protein level following TR compound treatment (Fig. 3D). Heat shock protein 60 (HSP60) is mitochondrial matrix chaperone required for maintaining the folding of imported proteins and HSP60 inhibitors are being investigated as potential anti-cancer compounds(51). Immunoblotting for HSP60 protein after TR-107 treatment of MDA-MB-231 cells showed that HSP60 levels were significantly reduced in a time-dependent manner (Fig. 3E). Together, these results demonstrate significant effects of TR-107 on mitochondrial proteins involved in energetic and homeostatic processes in a time-, dose- and ClpP-dependent manner.

### TR-107 inhibits OXPHOS in MDA-MB-231 cells

Because previous studies showed inhibition of OXPHOS by ONC201(25,27) and ADEP(52), we examined the effects of TR-107 on mitochondrial oxygen consumption and respiration using a Seahorse XF Analyzer. Incubation of MDA-MB-231 cells with TR-107 (25, 50 nM), resulted in a dose-dependent decline in oxygen consumption rate (OCR) (Fig. 4A). By contrast, this was not observed in ClpP-KO MDA-MB-231 cells (Fig. 4B). We observed significant dose-dependent decline in basal mitochondrial respiration, ATP production-linked oxygen consumption, maximal respiration, and spare respiratory capacity following TR-107 treatment in WT but not ClpP-KO cells (Fig. 4C). Correspondingly, TR107 incubation caused a dose-dependent increase in extracellular acidification (ECAR, Fig. 4C). Taken together, our results demonstrate substantial disruption of OXPHOS by TR107 in a dose and ClpP-dependent manner.

**Figure 4.**
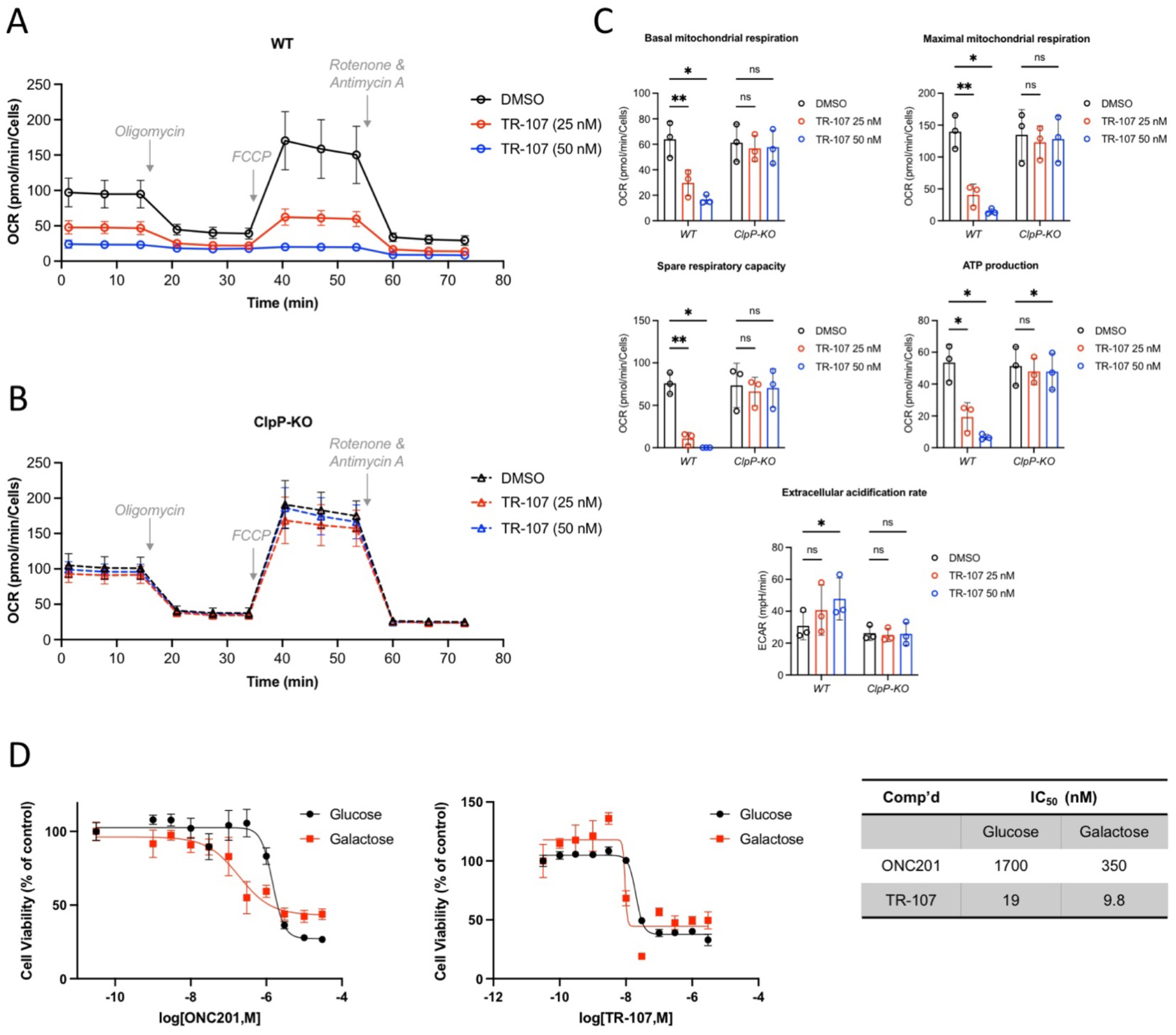
TR-107 reduces mitochondrial metabolic functions in MDA-MB-231 cells. Oxygen consumption rate (OCR) of MDA-MB-231 cells treated with DMSO or TR-107 at indicated concentrations for 24 hours was measured by Seahorse XF Analyzer as described in Materials and Methods. Trace of oxygen consumption data following TR-107 treatment in (A) WT and (B) ClpP-KO MDA-MB-231 cells, representative of N=3 experiments. Indicated compounds were added at concentrations described in Materials and Methods. C) Bar charts of data shown in (A) and (B), values represent mean ± SEM, N=3. P-value < 0.05 (*), 0.001 (**). D) Cell viability assays (Hoechst stain, 72 hours) of MDA-MB-231 cells treated with ONC201 (left) or TR-107 (right) at indicated concentrations in media containing either glucose (black) or galactose (red). Values represent mean ± SEM, representative of N=3. Average IC_50_ values are presented in table (right).

As shown by Greer *et al*., ONC201 treatment resulted in a greater dependence on glycolysis for cell survival and switching media carbon sources from glucose to galactose enhanced cell growth inhibition by ONC201(25). We similarly compared the effects of incubating cells in galactose instead of glucose on cell growth inhibition by TR-107. As expected for ONC201, galactose shifted the dose response curve for growth inhibition by ~5-fold (IC_50_ = 1.7 μM (glucose) and 350 nM (galactose)). While the effects on TR-107 on growth inhibition after galactose substitution were much less marked (~2-fold change; IC50 = 19 nM (glucose) and 9.8 nM (galactose)) (Fig. 4D), these results are consistent with OXPHOS inactivation and an increased reliance on glycolysis after TR-107 treatment.

### Pharmacokinetic properties of TR-107

With TR-107 and other leading TR compounds showing significantly enhanced potency over ONC201 and functional effects indicative of specific ClpP activation, we compared the PK properties of a set of TR compounds, ONC201 and ONC212/TR-31. The PK studies were conducted by tail vein injection or oral gavage in ICR mice as described in Materials and Methods. Based on the enhanced potency of this set of TR compounds we anticipated an effective oral dose to be ~10 mg/kg or less in a murine model of breast cancer. Therefore, we chose to evaluate the oral exposure of these agents with a single dose of 10 mg/kg and in many cases a 2 mg/kg i.v dose allowing for a F% determination. Importantly, the results of our studies showed significant differences in PK properties among these compounds (Table 1).

**Table 1.** Pharmacokinetic analysis of select compounds in mice. Each arm of pharmacokinetic were performed as described in Materials and Methods. Blood collection for ONC201 at 25 mg/kg (oral) was 0.083-4 hours. The TR compounds were administered intravenously (“i.v.”) (2.0 mg/kg) or by oral gavage (“oral”) (10 mg/kg) in vehicle, described in Materials and Methods. *A 2.5X increase in oral gavage dose gave a 46X increase in exposure (AUC). Three male ICR mice were utilized per study arm.

| Comp'd | Admin (mg/kg) | T<sub>1/2</sub> (hr) | C<sub>max</sub> (ng/mL)<br>[μM] | AUC (0- ∞;<br>h*ng/mL) | F% |
| --- | --- | --- | --- | --- | --- |
| ONC201 | 2.0, i.v. | 0.26 | 122 [0.31] | 50.7 | N/A |
|  | 10, oral | 0.31 | 8.99 [0.023] | 3.13 (1X) | 1.2 |
|  | 25, oral | --- | 195 [0.50] | 145 (46X)* | N/A |
| TR-27 | 2.0, i.v. | 1.06 | 154 [0.35] | 155 | N/A |
|  | 10, oral | 1.16 | 127 [0.29] | 255 | 33 |
| ONC212/TR-31 | 2.0, i.v. | 1.68 | 950 [2.2] | 638 | N/A |
|  | 10, oral | 1.49 | 282 [0.64] | 449 | 14 |
| TR-65 | 2.0, i.v. | 0.73 | 636 [1.5] | 351 | N/A |
|  | 10, oral | 0.95 | 294 [0.67] | 584 | 33 |
| TR-57 | 2.0, i.v. | 1.52 | 1240 [3.0] | 886 | N/A |
|  | 10, oral | 1.40 | 1710 [4.1] | 2710 | 61 |
| TR-107 | 10, oral | 0.90 | 1440 [3.7] | 2360 | N/A |

Of the impridone analogs closely related to ONC201, TR-27, ONC212/TR-31 and TR-65 showed enhanced exposure over ONC201, increased F%, and half-lives ranging from ~0.95-1.68 hours. Our initial studies of ONC201 using either an intravenous dose of 2 mg/kg or oral dose of 10 mg/kg showed markedly different results than what has been previously reported for higher doses. Both arms of our study showed a short t_1/2_ (0.26-0.31 hours) and poor oral bioavailability (F% = 1.2%) vs. a reported t_1/2_ of 6.4 hours in mouse(18). Notably, a 25 mg/kg oral dose of ONC201 showed a 46X increase in exposure (AUC) vs the 10 mg/kg study. In addition, ONC212/TR-31 displayed a terminal half-life of 1.49 hours (10 mg/kg) whereas the reported t_1/2_ of an oral dose of 125 mg/kg was 4.3 hours(53) (Table 1).

TR-57 and TR-107 displayed the highest exposure (AUC) of the tested agents when administered at 10 mg/kg orally (2710 and 2360 hr*ng/mL respectively, Table 1). Both compounds showed rapid absorption via oral administration with an F% of 61% (TR-57). In addition, protein binding studies for TR-107 shows a serum free fraction of 10% (mouse, Fig.S3C). Because of the favorable PK characteristics observed with TR-107 and its efficient molecular design, this compound was selected for advancement into animal xenograft studies described below.

### TR-107 inhibits tumor growth in a MD-MBA-231 mouse xenograft model

To determine the *in vivo* anti-tumor efficacy of TR-107, we examined dose-dependent responses using an MDA-MB-231 mouse xenograft model. MDA-MB-231 cells in Matrigel were orthotopically injected into the flank mammary fat pad of female NCr nu/nu mice (10/group) as described in Materials and Methods and Supplemental Information. Vehicle control (Group 1) and TR-107 treatments (4 mg/kg (Group 2), 8 mg/kg (Group 3)) were administered by oral gavage at frequencies described in Table S1.

The results of these studies demonstrated a dose- and time-dependent reduction in tumor volume. The mean tumor volume decreased for both treatment groups (G2, G3) with ~50% reduction in tumor volume observed at day 26 (Fig. 5A). Comparing the two different groups over the course of the study did not reveal significant advantages of one dosing regimen over the other in tumor volume reduction (Fig. 5A, 5B) or survival (Fig. 5C). A ~35% increase in median survival was also observed in mice treated with TR-107 for both groups (Fig. 5C). Consistent with ClpP activation, analysis of the tumor lysates from control and treated animals (G3), showed a substantial loss of mitochondrial DNA (Fig. 5D). TR-107 treatment was well tolerated and less than 5% weight loss was observed even at the highest dosing regimen (Fig. S3). Taken together, TR-107 showed significant efficacy in prevention of tumor growth in the MDA-MB-231 mouse xenograft model.

**Figure 5.**
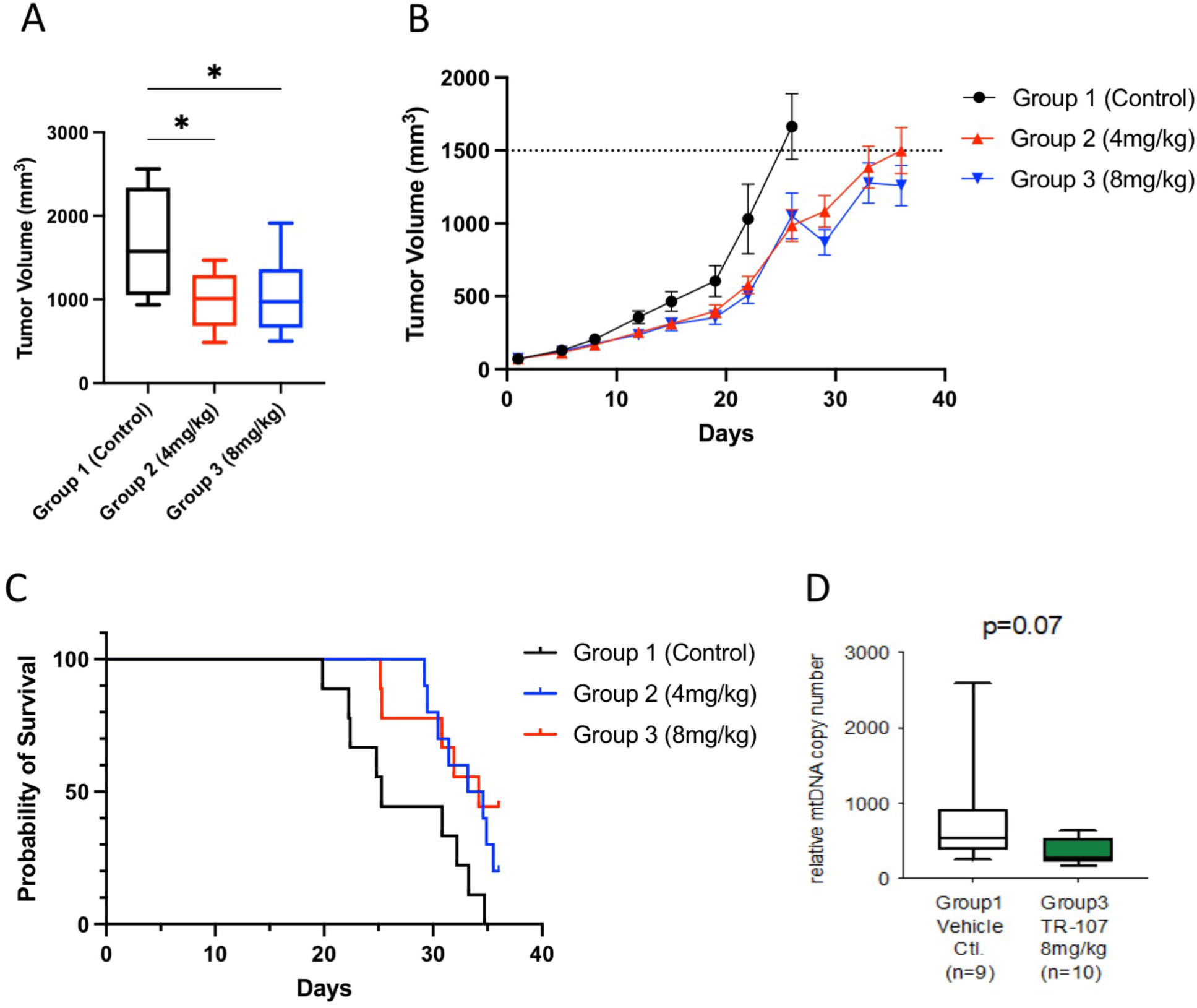
TR-107 prevents tumor growth in MDA-MB-231 xenograft model. A) Box-and-whisker plot of tumor volume (mm^3^) of MDA-MB-231 xenografts following TR-107 treatment at indicated concentrations at Day 26 (Table S2). N=8 (Group 1) and 10 (Groups 2, 3). P-value <0.05 (*) B) Average tumor volume following TR-107 administration as described in Materials and Methods. The tumor volume threshold (1500 mm^3^) is indicated by dashed line. N=10 per group. Values represent mean ± SEM. C) Kaplan-Meier graph of mouse survival following TR-107 administration as described in Materials and Methods and Table S2. Long-rank (Mantel-Cox) p-value of 0.0163. Individual mice were recorded as leaving the study following death, tumor volume exceeding threshold (1500 mm^3^), or end of study period. D) Box-and-whisker plot of relative mitochondrial DNA (mtDNA) copy number present in mouse xenograft lysate following conclusion of mouse study. N=9 (Group 1) and N=10 (Group 3).

## Discussion

ONC201, a screening hit with previously described CNS activity(22), was the first imipridone molecule shown to be effective against a variety of cancer models(17,19,22,50,54–56). Chemically similar derivatives of ONC201 (ONC206 and ONC212/TR-31) were recently identified and both ONC201 and ONC206 are under evaluation in clinical trials. Despite evidence suggesting that ONC201 acts through effects on the dopamine receptor(56), the discovery of the mitochondrial protease ClpP as the target for ONC201 and chemically related compounds(26,27,57) has redefined our understanding of the anticancer mechanism of these agents. Moreover, the identification of ClpP as the major target has enabled the design and characterization of highly potent and specific ClpP activators. In this study, we describe the advancement of the chemically diverse TR compounds as selective ClpP activators, which includes molecules of different chemical scaffolds from which agents for preclinical assessment were selected. We characterize one of these compounds, TR-107, for effects on mitochondrial metabolism and protein turnover, PK properties and TNBC growth inhibition both *in vitro* and *in vivo*. Importantly, these studies further demonstrate the pivotal role for ClpP activation in the mechanism of TR-107 action.

From the collection of TR compounds, we were able to perform initial structure activity relationship (SAR) studies using cell growth inhibition and ClpP activation as determinants. Nanjing Gator Meditech was the first to accurately determine that for the imipridone scaffold, substitution at the V- and Z-positions by small lipophilic moieties was preferential to substitution at the W-position, yielding nM activity in cancer cells(58). Our survey of residues for the other benzyl ring determined that the nitrile residue at position Y- was preferred. The combination of these two optimized benzyl residues on the imipridone scaffold (TR-65) resulted in markedly improved potency with regards to both ClpP activation and growth inhibition(26) compared to ONC201 and ONC206.

Two key chemical features of the imipridone chemical core are a basic nitrogen of the 3-piperidine moiety and an imbedded acylguanidine. The importance of the basic nitrogen of the piperideine residue to activity is demonstrated through substitution of the nitrogen atom with carbon atom (e.g. X position, TR-109 and TR-122, Fig. 1A). Both TR-109 and TR-122 are significantly less potent than TR-65 displaying μM IC_50_ values for TNBC cells. We previously reported the modification of the acylguanidine moiety to prepare TR-57 and the homologated analogs TR-79 and TR-80 for target identification studies(26). We now report that further modification of TR-57 removing the carbonyl oxygen and methyl residue (R group) gives TR-107, a compound of equal potency. In addition, expansion of the 5-membered ring of the imipridone scaffold by one methylene unit yields TR-98, a compound that is equipotent to TR-65. In summary, the additional modifications described here resulted in two of the most potent ClpP activators (TR-98 and TR-107) so far reported and further support that the imipridone chemical scaffold is not a requirement of potent ClpP activation.

Because of high potency in cell growth and ClpP activation assays, TR-107 was selected for further mechanistic, pharmacokinetic and animal model studies. Other attributes include a low molecular weight (<400 amu), highly potent and defined ClpP dependence, excellent PK properties and efficacy against TNBC models both *in vivo* and *in vitro*. Using two independent ClpP-KO cell lines, our studies confirm the mitochondrial protease ClpP as the key target for TR-107 and related compounds. This includes effects on cell growth, OCR, and mitochondrial protein level as determined by immunoblotting. While ONC201, at concentrations far higher than that is necessary to activate ClpP or inhibit cell growth, has shown effects on dopamine receptors D2/D3(56,59), our data(26) and that of others(25,27) strongly argues against a major role of the dopamine receptors in growth inhibition.

In strong support of a mitochondrial mechanism of action, we observed TR-107 treatment results in reduction of multiple mitochondrial proteins. Some of the most significantly affected proteins, in addition to TFAM and TUFM, included ACO2 and IDH2, which catalyze essential reactions in the TCA cycle. Similarly, respiratory chain proteins NDUFS3 and SDHA declined strongly. Lastly, we observed the ClpP-dependent loss of other essential mitochondrial proteins including POLRMT and HSP60, both of which are predicted to have significant effects on mitochondrial function through regulation of mitochondrial gene replication and protein chaperone functions. Both POLRMT and HSP60 have been shown to be viable drug targets for cancer research(10,51), and provide potential further explanation for the anticancer properties of these compounds. While this is a partial list of all mitochondrial proteins observed to decline following TR-107 treatment (E. Fennell, unpublished), based on these results we propose that inactivation of key mitochondrial functions is, in part, responsible for the growth inhibition observed in these studies.

Mitochondrial dysfunction and metabolic reprogramming are accepted hallmarks of cancer(6,60–63). Inhibiting mitochondrial processes such as OXPHOS has gained interest as an alternative approach to combatting cancer(13,64–69). Our studies confirm the original findings of Greer *et al*. that ONC201 targets mitochondrial metabolism. In addition to TCA cycle enzymes, we observed proteins with essential roles in OXPHOS that were reduced after ONC201 and TR-57 exposure. The inhibition of OXPHOS was confirmed by mitochondrial respiration analysis and galactose experiments. Interestingly, substituting media glucose for galactose had a much greater effect on the IC50 values of ONC201 compared to TR-107. These data further illustrated the superior potency of TR-107 and suggests that the effects of TR-107 may be less dependent of nutritional conditions (e.g. glucose availability) compared to ONC201. Taken together, the results of our studies demonstrate an increased reliance on glycolysis as compensatory response to inhibition of mitochondrial metabolism by ClpP activation.

Previous attempts to inhibit OXPHOS or specific enzymes in the TCA cycle have met with limited success for cancer treatment. The repurposing of metformin [1,1-dimethylbiguanide] as an anticancer agent has been investigated in both research and clinical settings(65,69,70). As a weak inhibitor of mitochondrial complex I, metformin appears to lack the potency and specificity to effectively treat cancer as a single agent or in combination via complex I inhibition. The complex I inhibitor, BAY 87-2243 is a highly potent inhibitor of complex I with a more complex chemical structure than metformin featuring lipophilic pharmacophores absent in metformin. Notably, clinical evaluation of BAY 87-2243 was terminated after treatment of patients (NCT01297530). A focused contemporary approach by Stuani *et al*. to target mutant forms of IDH 1 and 2 and thereby reducing the oncometabolite (R)-2-hydroxyglutarate (2-HG), led to agents that showed initial efficacy but lost their effectiveness with established disease(71).

As single agents, we observed that ONC201 and the TR compounds induced a growth stasis response in these TNBC models. Even at the highest concentrations tested, little evidence for apoptosis was observed. In this way our results are consistent with earlier studies in TNBC that showed a similar absence of apoptosis after treatment with ONC201(21,25). Recent studies suggest that addition of TRAIL or TRAIL receptor agonists induces an apoptotic response in cells that have previously displayed a growth stasis response to ONC201(72–74). Moreover, combination of imipridones with BCL-2 inhibitors has shown promise for increased cancer cell apoptosis(23,75). Given the greatly improved potency of the TR compounds, it will be important to determine if low concentrations of TR compounds more effectively sensitize cells to TRAIL treatment or synergize with TRAIL ligand or other agents to increase apoptosis(76).

Lastly, the development of potent and selective ClpP agonists addresses a key limitation for ONC201, ONC206, and ONC212/TR-31 series of compounds regarding the requirement of high doses (50-130 mg/kg) to achieve *in vivo* efficacy(53). The enhanced potency of ONC206 and TR-31/ONC212 in cellular growth assays did not result in lower *in vivo* efficacious doses. The pharmacokinetic data reported herein, coupled with previously reported results, points to a complex picture of pharmacokinetic behavior of ONC201 and ONC212/TR-31 at therapeutically relevant doses.

TR-107 at an oral dose of 10 mg/kg results in similar t1/2 and serum exposure (AUC) to an equivalent oral dose of TR-57. In addition to improved PK properties, TR-107 effectively reduced tumor volume and increased median survival *in vivo* at 4 and 8mg/kg dosages. Further analysis of xenografts revealed decrease of mitochondrial DNA following TR-107 administration, providing additional evidence for mitochondrial mechanism of action.

In summary, our studies show chemical optimization of ONC201 has resulted in novel ClpP activators with increased potency in both *in vitro* and *in vivo* studies. TR-107 showed excellent potency in TNBC cell growth inhibition and provide further evidence that ClpP activation inhibits growth through loss of mitochondrial metabolic functions. Our *in vivo* studies show that these compounds are highly orally bioavailable, well-tolerated, and effective at reducing tumor burden in an MDA-MB-231 murine xenograft model. In total, our observations support the clinical evaluation of TR-107 for the treatment of TNBC.

## Author Contributions

EMJF performed all cell viability and apoptosis assays, cell sample preparation and immunoblotting, and statistical analysis of these experiments, and manuscript writing. LJAC generated the ClpP-KO SUM159 cell line. JDW performed *in vitro* ClpP activity assays and assisted with sample preparation and immunoblotting. KDM performed Seahorse XF Analyzer experiments with assistance from EMJF. EL performed SPR experiments and Kd determination. YEG and SL provided MDA-MB-231 cell lines (WT and ClpP-KO). Quantification of mtDNA from mouse xenografts was performed by YEG. EJI designed TR compounds. HL, AAI, and DSK contributed to agent selection, study design of *in vivo* experiments, and data interpretation. PRG and EH contributed to study design. HA, CJD, WAH, and SL provided material support. LMG provided material support, manuscript writing, experimental design and coordination of these studies.

## Acknowledgements

The authors would like to acknowledge Dr. Michael P. East for invaluable scientific discussion and generous help with experimental troubleshooting. This project is supported by grants from National Institutes of Health to LMG (5R01GM138520-02) and CJD (R35CA232113). Additional research on this project is supported by a Canadian Institutes of Health Research Project grant (PJT-173345) and Canadian Cancer Society Innovation Grant (706282) to WAH. KDM was supported by NCI T32CA009156. This research was supported in part by the Intramural Research Program of the National Cancer Institute, Center of Cancer Research (ZIA SC 007263 to SL).

