## Supplemental Information for "Characterization of TR-107, a Novel Chemical Activator of the Human Mitochondrial Protease ClpP"

^1^Department of Pharmacology and the Lineberger Comprehensive Cancer Center, University of North Carolina at Chapel Hill, Chapel Hill, North Carolina. ^2^Department of Biochemistry, University of Toronto, Toronto, Ontario M5G 1M1, Canada. ^3^Women’s Malignancies Branch, Center for Cancer Research, National Cancer Institute, National Institutes of Health, Bethesda, Maryland. ^4^New York Presbyterian Brooklyn Methodist Hospital, Department of Radiation Oncology, Brooklyn, New York. ^5^Madera Therapeutics LLC, Chapel Hill, North Carolina. ^6^Institute of Theoretical and Experimental Biophysics, Russian Academy of Sciences, Pushchino, Russian Federation, 142292. ^7^Department of Chemistry, University of Toronto, Toronto, Ontario M5S 3H6, Canada.

Supplemental Figures and Figure Legends

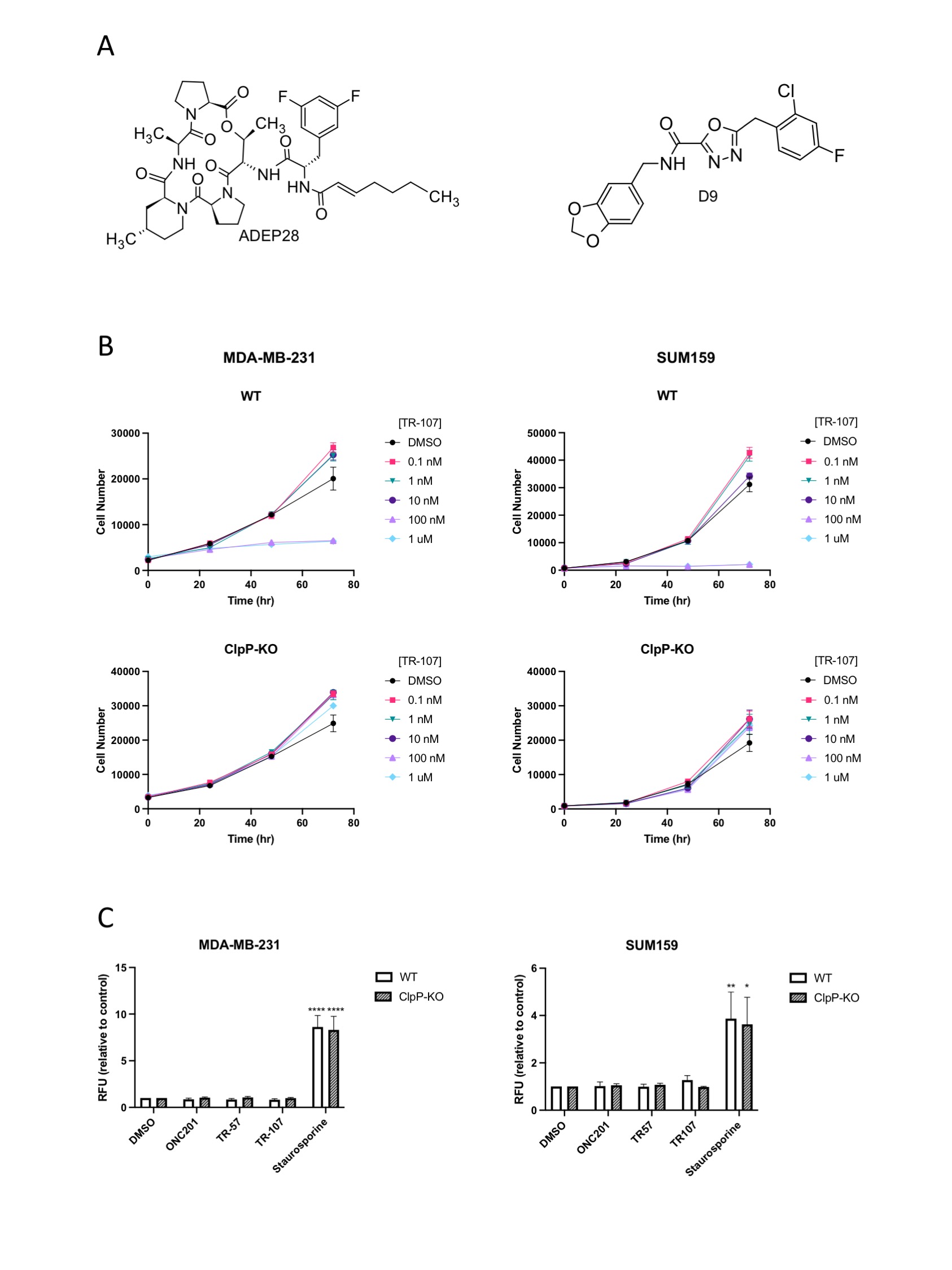

**Figure S1**. **TR compounds do not induce cell death in breast cancer cell lines.** A) Chemical structures of ClpP activators ADEP28 and D9. B) Total cell count assay in SUM159 and MDA-MB-231 cells (WT and ClpP-KO). Cells were treated with indicated drug concentrations for 0, 24, 48, and 72 hours and counted using Hoechst stain. Values represent mean cell count ± SEM, Representative of N=2. C) *In vitro* caspase-3/7 activity assay in SUM159 and MDA-MB-231 cells (WT and ClpP-KO). Cells were treated with 10 µM ONC201, 150 nM TR-57, 100 nM TR-107 or 100 nM staurosporine for 24 hours and caspase activity was measured via fluorometric caspase activity assay. Values represent mean fluorescence value ± SEM, N=3, p-value < 0.05 (*), 0.01(**), 0.0001 (****)

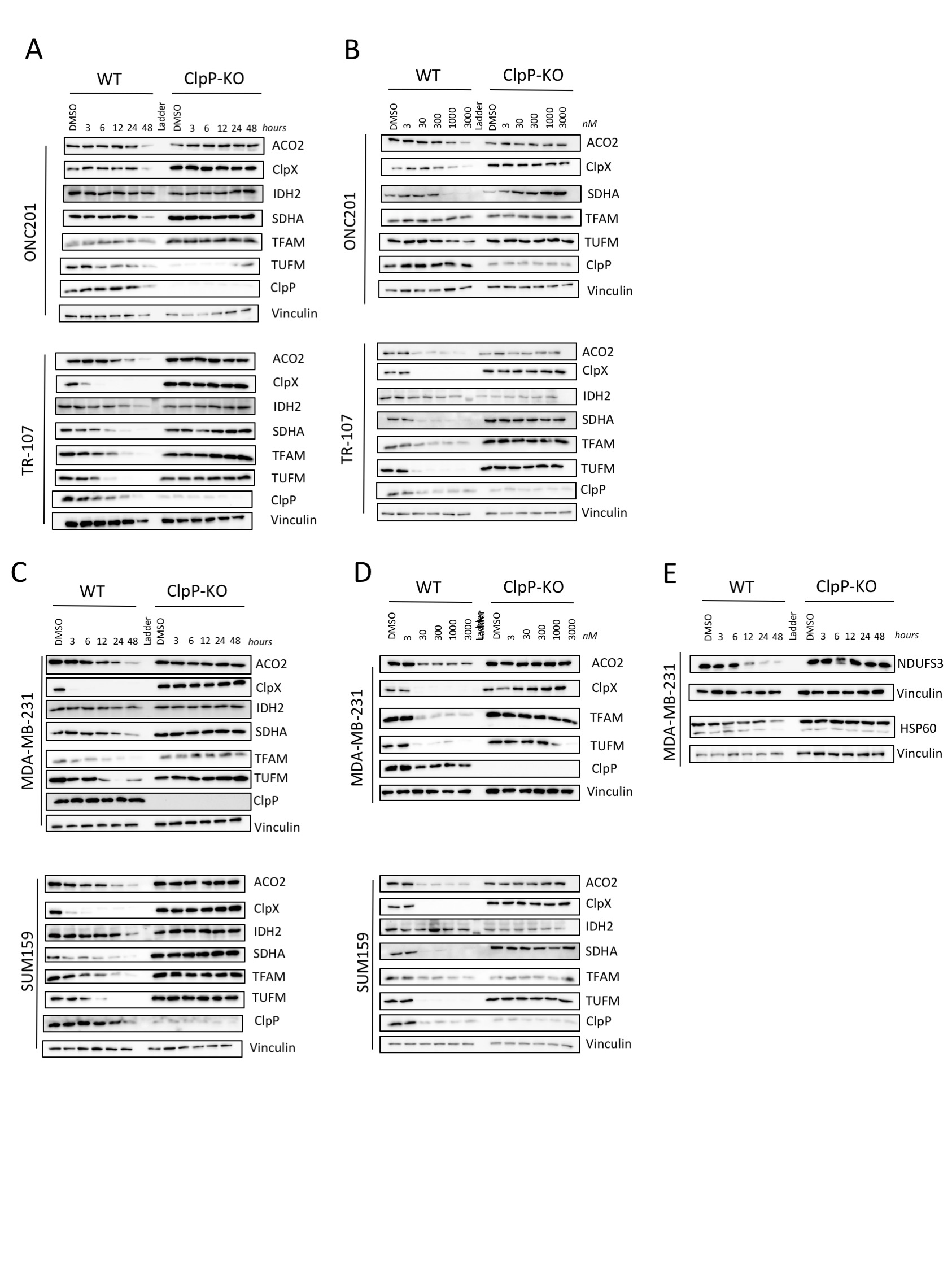

**Figure S2. ONC201, TR-107 and TR-57 induce loss of mitochondrial proteins in SUM159 cells in a ClpP-dependent manner.** SUM159 cells were treated with (A) 10 μM ONC201 or 100 nM TR-107 for indicated time points or (B) indicated doses of ONC201 or TR-107 for 24 hours. SUM159 and MDA-MB-231 cells were treated with (C) 150 nM TR-57 for indicated timepoints or (D) indicated doses of TR-57 for 24 hours and immunoblotted for various mitochondrial proteins as described in Methods. (E) MDA-MB-231 cells were treated with 150 nM TR-57 for indicated timepoints and immunoblotted for NDUFS3 and HSP60. Data representative of N=3 (A-D) or N=2 (E) experiments.

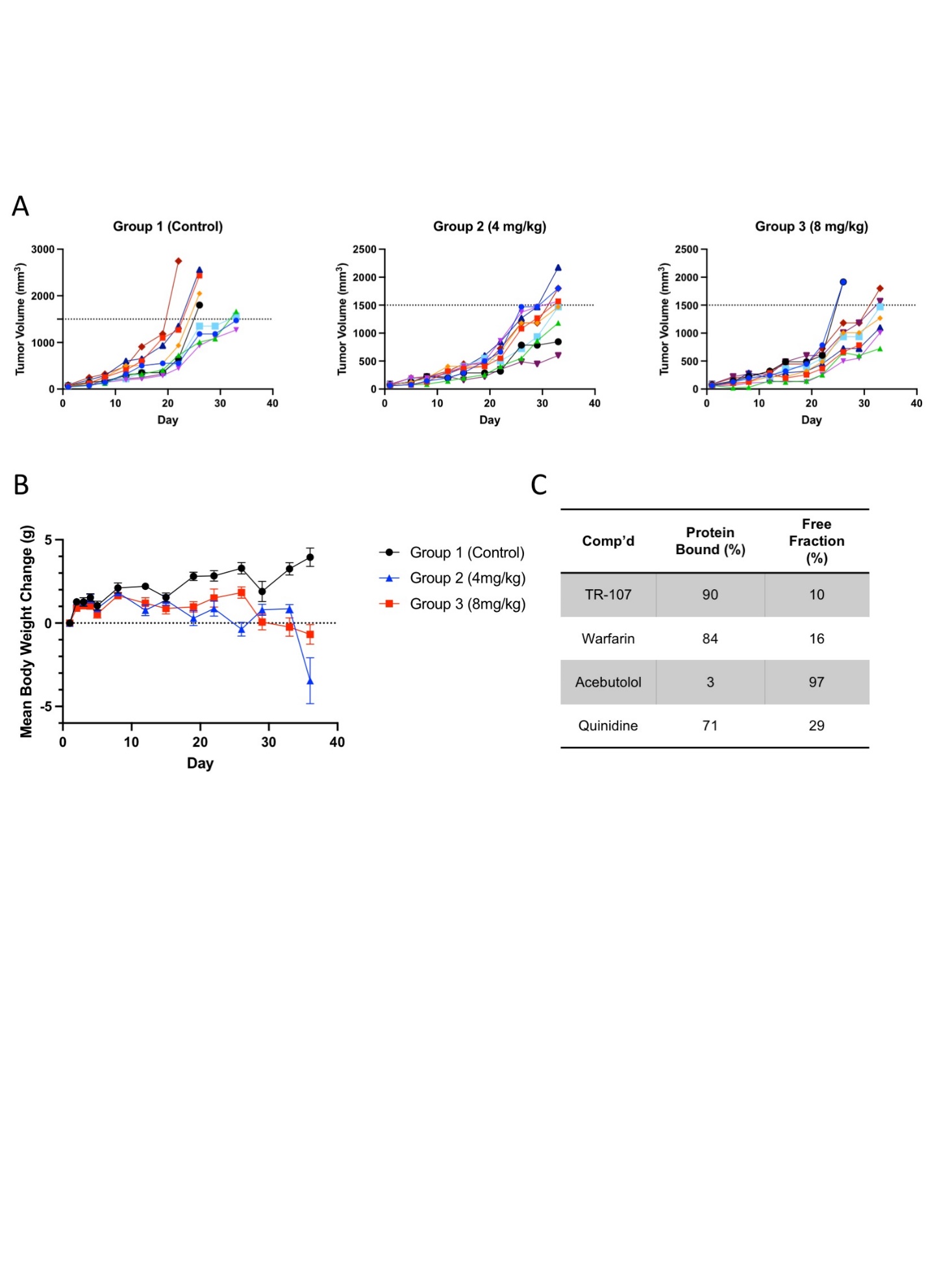

**Figure S3. TR-107 inhibits tumor growth in MDA-MB-231 mouse xenograft model.** A) Graphs of individual tumor volume measurements at indicated time points. Each color represents a single mouse participating in the study, N=10 per group. B) Average change in body weight compared to initial measurement of mice receiving vehicle (Group 1) or TR-107 treatment (Group 2 (4 mg/kg) or Group 3 (8 mg/kg)) at indicated time points. Values represent average ± SEM. N=10 per group. C) Murine plasma protein binding profile of TR-107 and control compounds (Warfarin, acebutolol, and quinidine). Values represent mean of N=2 replicates.

**Table S1. TR-107 dosing and treatment protocol for murine MDA-MB-231 xenograft study.** All compounds were dissolved in vehicle as described in Methods and were administered orally (P.O.) to 10 mice per group on schedules indicated.

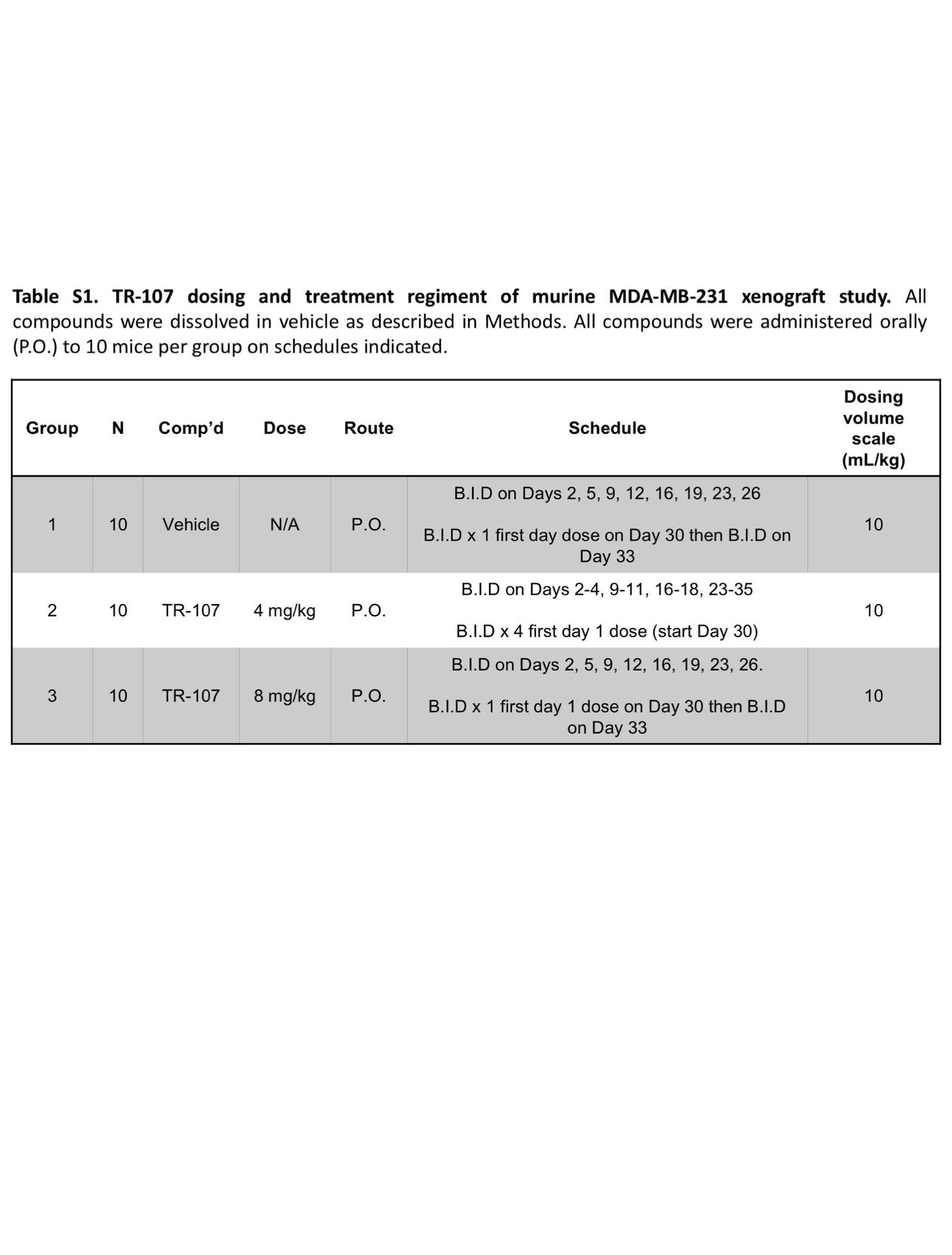

Supplemental Methods

**Mouse Xenograft Studies**

**Mice**

Female athymic nude mice (Crl:NU(Ncr)-Foxn1*^nu^,* Charles River) were eight weeks old with a body weight (BW) range of 19.0 to 27.0 grams on Day 1 of the study. Animals were fed ad libitum water (reverse osmosis, 1ppm Cl) and NIH 31 Modified and Irradiated Lab Diet® consisting of 18.0% crude protein, 5.0% crude fat, and 5.0% crude fiber. Mice were housed on irradiated Enrich-o’cobs^TM^ Laboratory Animal Bedding in static microisolators on a 14-hour light cycle at 20-22 C and 40-60% humidity. Charles River Discovery Services North Carolina (CR Discovery Services) specifically complies with the recommendations of the Guide for Care and Use of Laboratory Animals with respect to restraint, husbandry, surgical procedures, feed and fluid regulation, and veterinary care. Animal care and use program at CR Discovery Services is accredited by the Association for Assessment and Accreditation of Laboratory Animal Care International.

**Tumor Cell Culture**

MDA-MB-231 human breast adenocarcinoma cells were grown to mid-log phase in RPMI-1640 supplemented with 10% FBS, 2 mM glutamine, 100 units/mL sodium penicillin G, 25 µg/mL gentamicin, 100 µg/mL streptomycin sulfate, 0.075% sodium bicarbonate, 10 mM HEPES, and 1 mM sodium pyruvate. Cells were cultures in 37°C and 5% CO_2_.

***In Vivo* Implantation and Tumor Measurement**

MDA-MB-231 cells were harvested during exponential growth and resuspended in cold, sterile DPBS. Each animal received an orthotopic injection of 5x10^6^ cells (0.05 mL volume) into the mammary fat pad. Tumor growth was monitored as the average tumor size approached target range of 60-100 mm^3^. Tumors were measured in two dimensions via caliper and volume was calculated using the following formula:

$$Tumor Volume= \frac{w^{2} x l}{2}$$

Where *w* =width and *l*= length (mm) of the tumor. Tumor weight was estimated based on the assumption that 1 mg = 1 mm^3^. Fourteen days later (Day 1), animal were sorted into groups (n=10 per group) with a mean tumor volume of 70-73 mm^3^.

**Treatment**

Mice began dosing on Day 1 as summarized in Table S1. Compounds were administered orally (p.o.) in a dosing volume of 10 mL/kg. Twice daily doses (B.I.D) were administered at least 6 hours apart.

**Endpoint and Tumor Growth Delay (TGD) Analysis**

Tumors were measured using calipers biweekly and animals were euthanized upon reaching tumor volume >1500 mm^3^ or on the final study day (Day 36), whichever came first. Time to endpoint (TTE) for analysis was calculated for animals that exited the study due to tumor volume using the following equation:

$$TTE= \frac{{log}_{10}\left( endpoint volume \right)-b}{m}$$

Where TTE is expressed in days, endpoint volume is expressed in mm^3^, *b* is the y-intercept, and *m* is the slope of the line obtained by linear regression of a log-transformed tumor growth data set. This dataset consisted of the first observation that exceeded 1500 mm^3^ and the preceding three consecutive measurements. Animals that did not reach the tumor volume threshold were assigned a TTE value of 36 days (equivalent to study end). In instanced where this equation yields a TTE day preceding the day prior to reaching endpoint or exceeded the day of reaching endpoint, a linear interpolation was performed to approximate TTE. Animals documented as having died of non-treatment-related causes due to accident or unknown etiology were excluded from TTE calculations and all further analysis. Animals documented having died due to treatment-related deaths or non-treatment-related due to metastasis were assigned a TTE value equivalent to day of death. Treatment outcome was evaluated from tumor growth delay (TGD) defined as the increase in the median TTE. In the treatment group compared to the control group.

**Sampling**

On Day 36, all animals in all groups were sampled. Tumors were excised, divided into two parts, and weighed. Part 1 was preserved in 5 mL fixative solution (4% formaldehyde, 2% glutaraldehyde in 0.1M cacodylate buffer) and stored at 4°C. Part 2 was trimmed to be less than 0.5 cm in at least one dimension, submerged in 5 volumes of RNAlater solution, and stored at -20°C.

**Statistical and Graphical Analyses**

GraphPad Prism 9.02 was used for all statistical analysis and graphical presentations. Study groups experiencing toxicity beyond acceptable limits (>20% group mean BW loss or >10% treatment-related deaths) or having fewer than five evaluable observations were not included in statistical analysis. Survival was analyzed by Kaplan-Meier method and logrank (Mantel-Cox) test was used to determine significance between overall survival experiences based on TTE values.

**Immunoblotting**

Primary and secondary antibodies used in immunoblot analysis are listed below:

| Name | Manufacturer | Catalog Number |
| --- | --- | --- |
| Anti-SDHA | Cell Signaling Technologies | 5839 |
| Anti-ClpP |  | 14181 |
| Anti-ACO2 |  | 6922 |
| Anti-TUFM | Invitrogen | PA5-27511 |
| Anti-IDH2 |  | PA5-79436 |
| Anti-POLRMT |  | PA5-28196 |
| Anti-ClpX | Abcam | 168338 |
| Anti-NDUFS3 |  | 183733 |
| Anti-HSP60 | BD Transduction Laboratories | 611562 |
| Anti-mtTFA (TFAM) | Santa Cruz Biotechnologies | sc-376672 |
| Anti-Vinculin |  | sc-73614 |
| Anti-Rabbit IgG HRP conjugate | Promega | W401B |
| Anti-Mouse IgG HRP conjugate |  | W402B |
